# Neurons in the inferior temporal cortex of macaque monkeys are sensitive to multiple surface features from natural objects

**DOI:** 10.1101/086157

**Authors:** Hiroshi Tamura, Haruki Otsuka, Yukako Yamane

**Affiliations:** Graduate School of Frontier Biosciences, Osaka University, Suita, Osaka 565-0871, Japan; Center for Information and Neural Networks, Suita, Osaka 565-0871, Japan

**Keywords:** inferior temporal cortex, material perception, object recognition, primary visual cortex, V4

## Abstract

Object surfaces contain a variety of visual features that help us to recognize them. To understand how this information is represented and processed in the brain, we prepared a set of images from natural object surfaces that maintained surface features but lacked contours. We examined spiking responses of neurons in the inferior temporal (IT) cortex of monkeys, which is a crucial structure needed for visual object recognition. About half of IT neurons responded to surface images with sharp selectivity, indicating that a significant fraction of these neurons contribute to object surface representation in a sparse manner. Responses of IT neurons were susceptible to image manipulations, including color removal, removal of luminance contrasts, and spatial structure degradation. This shows that multiple features are required for IT responses to surface images. Comparing neuronal response properties among IT, visual area 4 (V4), and primary visual cortex (V1) revealed properties of IT neurons that differed from those in the other visual processing regions. Additionally, some neuronal response properties were similar between IT and V4, but differed from those in V1, indicating that responses of IT neurons to surface images are constructed by hierarchical processing throughout the ventral visual pathway.

---

Surfaces of natural objects contain a variety of visual features, such as colors, luminance contrasts, and spatial patterns. Combinations of these features confer a unique visual appearance to object surfaces and help us to recognize them. Psychophysical and behavioral studies have shown that visual features of surfaces contribute to object recognition in humans and monkeys (Livingstone and Hubel 1987; Price and Humphreys 1988; Cavanagh 1991; Vogels 1999; Coss and Ramakrishnan 2000; Adelson 2001; Regan et al. 2001; Tanaka et al. 2001; Waitt et al. 2003; Rossion and Pourtois 2004; Changizi et al. 2006; Fleming et al. 2015).

In macaque monkeys, visual object recognition depends on the neural structures of the ventral visual cortical pathway (Mishkin et al. 1982), at the end of which is the inferior temporal (IT) cortex. The majority of IT neurons are generally believed to be sensitive to stimulus shape (Gross et al. 1972; Desimone et al. 1984; Tanaka et al. 1991). However, some neurons in IT are sensitive to surface-related features, such as colors (Desimone et al. 1984; Tanaka et al. 1991; Komatsu et al. 1992; Komatsu and Ideura 1993; Tamura and Tanaka 2001; Edwards et al. 2003; Conway et al. 2007; Harada et al. 2009), textures (homogeneous spatial patterns; Desimone et al. 1984; Tanaka et al. 1991; Komatsu and Ideura 1993; Wang et al. 2003; Köteles et al. 2008;), structured spatial patterns (inhomogeneous spatial patterns; Desimone et al. 1984; Tanaka et al. 1991), or three-dimensional structures defined by luminance gradient (Yamane et al. 2008). These IT neurons may represent a single visual feature and be insensitive to others. Human functional magnetic resonance imaging (fMRI) studies have found that texture and colors activate separate regions (Cant et al. 2009; Cavina-Pratesi et al. 2010), suggesting that individual IT neurons can represent only a single visual feature. Additionally, other surface-responsive neurons are sensitive to multiple visual features, although the incidence of these neurons has been unknown or small (Desimone et al. 1984; Tanaka et al. 1991; Komatsu and Ideura 1993).

Information derived from the surfaces of natural objects is transferred to IT cortex through a series of cortical regions, beginning at the primary visual cortex (V1), and continuing through visual area 2 (V2) and visual area 4 (V4) before reaching IT (see Kravitz et al. 2013, for a review). Neurons in these preceding regions are also sensitive to colors (Hubel and Wiesel 1968; Zeki 1973; Livingstone and Hubel 1984; Lennie et al. 1990; Schein and Desimone 1990; Tamura et al. 1996; Johnson et al. 2001; Tootell et al. 2004; Kusunoki et al. 2006; Kotake et al. 2009; Bushnell et al. 2011), textures (Knierim and Van Essen 1992; Arcizet et al. 2008; Freeman et al. 2013; Okazawa et al. 2015), structured spatial patterns (Gallant et al. 1996), and three-dimensional structures defined by luminance gradients (Hanazawa and Komatsu 2001). An as yet unresolved issue is whether surface-sensitive IT neurons simply inherit response properties from these preceding regions, or if further processing in IT confers them with unique surface-related response properties.

The first aim of the present study was to understand how the surface visual features derived from natural objects are represented in IT cortex. Although neurons have been shown to respond to surface-related visual features, quantifying the incidence and stimulus selectivity of these neurons is important for understanding the representation of these visual features in IT cortex. To address this issue, we extracellularly recorded spiking responses of single IT neurons in macaque monkeys. We quantified the incidence and stimulus selectivity of responsive neurons using a set of images derived from the surfaces of natural objects. The second aim of the study was to clarify whether the responses of these neurons depended on the presence of multiple visual features, such as color, luminance contrast, and spatial structure. For this, we examined responses of IT neurons to images in which some of the visual features had been modified or removed. The third aim was to determine how the visual information derived from object surfaces is processed in the hierarchically organized ventral visual cortical pathway. In addition to IT, we therefore recorded spiking responses of single neurons in V4 and V1 to the same set of images.

## Materials and Methods

We recorded neuronal responses from six monkeys (*Macaca fuscata*; body weight, 5.9–8.6 kg; Monkeys A, B, C, D, E, and F). We shared these monkeys with other research projects. Because some of the recording sessions required stable recording for more than an hour, we used an analgesized and paralyzed preparation. Although analgesia/paralysis might have affected neuronal activity, any effect was likely immaterial given that stimulus selectivity of V1 and IT neurons recorded from analgesized/paralyzed monkeys has been shown to be similar to that of awake-behaving monkeys (Wurtz 1969; Tamura and Tanaka 2001). Because general experimental procedures were similar to those described previously (Tamura et al. 2014), the description here will be brief. All experiments were performed in accordance with the guidelines of the National Institute of Health (1996) and the Japan Neuroscience Society and were approved by the Osaka University Animal Experiment Committee.

### Initial preparatory surgery

We prepared monkeys for recordings during an aseptic surgery in which a head restraint was implanted and the lateral and occipital part of the skull over the recording region was covered with acrylic resin. We performed the surgery under full anesthesia via inhalation of 1%–3% isoflurane (Forane, Abbott Japan, Tokyo, Japan) in nitrous oxide (70% N_2_O, 30% O_2_) through an intratracheal cannula. We gave monkeys an antibiotic (Pentcilin, Toyama Chemical, Tokyo, Japan; 40 mg/kg, i.m.) and an anti-inflammatory and analgesic agent (Voltaren, Novartis, Tokyo, Japan; or Ketoprofen, Nissin Pharmaceutical, Yamagata, Japan) immediately after the surgery and continued during the first postoperative week. After 1–2 weeks of recovery, we examined the eyes to select appropriate contact lenses that allowed images placed 57 cm from the cornea to be focused on the retina. We took photographs of the retinal fundus to determine the position of the fovea.

### Animal preparation for neural recording

On the day of neural recording, we sedated monkeys using intramuscular injections of atropine sulfate (0.1 mg/kg) and ketamine hydrochloride (12 mg/kg). During preparation, we analgesized monkeys via inhalation of 1%–3% isoflurane in nitrous oxide (70% N_2_O, 30% O_2_) through an intratracheal cannula, and infused them with the opioid fentanyl citrate (Fentanest, Daiichi Sankyo; 0.035 mg/kg/h) in lactated Ringer’s solution. We drilled a small hole (~5 mm) in the resin-covered skull and made a small slit (2 mm) in the dura for electrode insertion. We dilated the pupil of the eye contralateral to the recording hemisphere and relaxed the monkeys’ lenses using 0.5% tropicamide/0.5% phenylephrine hydrochloride (Mydrin-P, Santen, Osaka, Japan). We then covered the cornea of the contralateral eye with a contact lens of appropriate refractive power and curvature, and with an artificial pupil (diameter, 3 mm) so that the eye would focus on images placed 57 cm away. After inserting the recording electrode, we added vecuronium bromide (Masculax, MSD, Tokyo, Japan; 0.06 mg/kg/h) to the infusion solution to prevent eye movements during the recordings of neuronal activity.

#### Neural recordings

We made multiple single-unit recordings from IT, V4, and V1 using a single-shaft electrode with 32 recording probes arranged linearly (A1X32-10 mm 50-413, A1X32-10 mm 100-413; NeuroNexus, Ann Arbor, MI, USA) or an eight-shaft electrode, with each shaft being a tetrode having four recording probes at the tip arranged in a rhombus (A8X1 tetrode-2 mm 200-312; NeuroNexus, Ann Arbor, MI, USA). For the eight-shaft electrode, the centers of adjacent shafts were separated by 0.2 mm. The distance between the centers of adjacent recording probes was 50 µm or 100 µm when using the single-shaft electrode, and 25 µm when using the eight-shaft electrode. Because we could observe spiking activity from the same neuron in two or more adjacent probes, we isolated single neuron activity offline using custom-made software (see Kaneko et al. 1999; Kaneko et al. 2007; Tamura et al. 2014, for details). After each recording session, we provided the monkeys analgesics and antibiotics and returned them to their home cages. Each recording session lasted up to 7 h, and we waited at least a week before the next recording session in a particular monkey.

The recording sites in IT cortex were located between the superior temporal sulcus and the anterior middle temporal sulcus, and anterior to the posterior middle temporal sulcus. Those in V4 were located between the superior temporal sulcus and the lunate sulcus, and those in V1 were located in the surface of the occipital cortex well behind the lunate sulcus.

### Visual stimuli

We prepared a stimulus set consisting of 64 images of natural objects (original images; Fig. 1A): stones (*n* = 8, *#1–8*), tree bark (*n* = 8, *#9–16*), leaves (*n* = 8, *#17–24*), flowers (*n* = 8, *#25–32*), fruits and vegetables (*n* = 8, *#33–40*), butterfly wings (*n* = 8, *#41–48*), feathers (*n* = 8, *#49–56*), and skins and furs (*n* = 8, *#57–64*); and a blank image (*#65*) that has the same pixel values as the background. Most images were the outer surface of objects but two (*#35* and *#36*) were cut surfaces. We selected objects to include a variety of colors, luminance contrasts and spatial structures. All objects were natural (i.e., not man-made), although the monkeys might never have seen some of them in their daily lives. The experimenters photographed the images in indoor and outdoor environments. During the photographing and processing, we enlarged some images and shrunk others, and therefore the images were not all in the same scale. From the original photographs, we cut out square images, removing outer contours. All images were 6° in visual angle.

**Figure 1.**
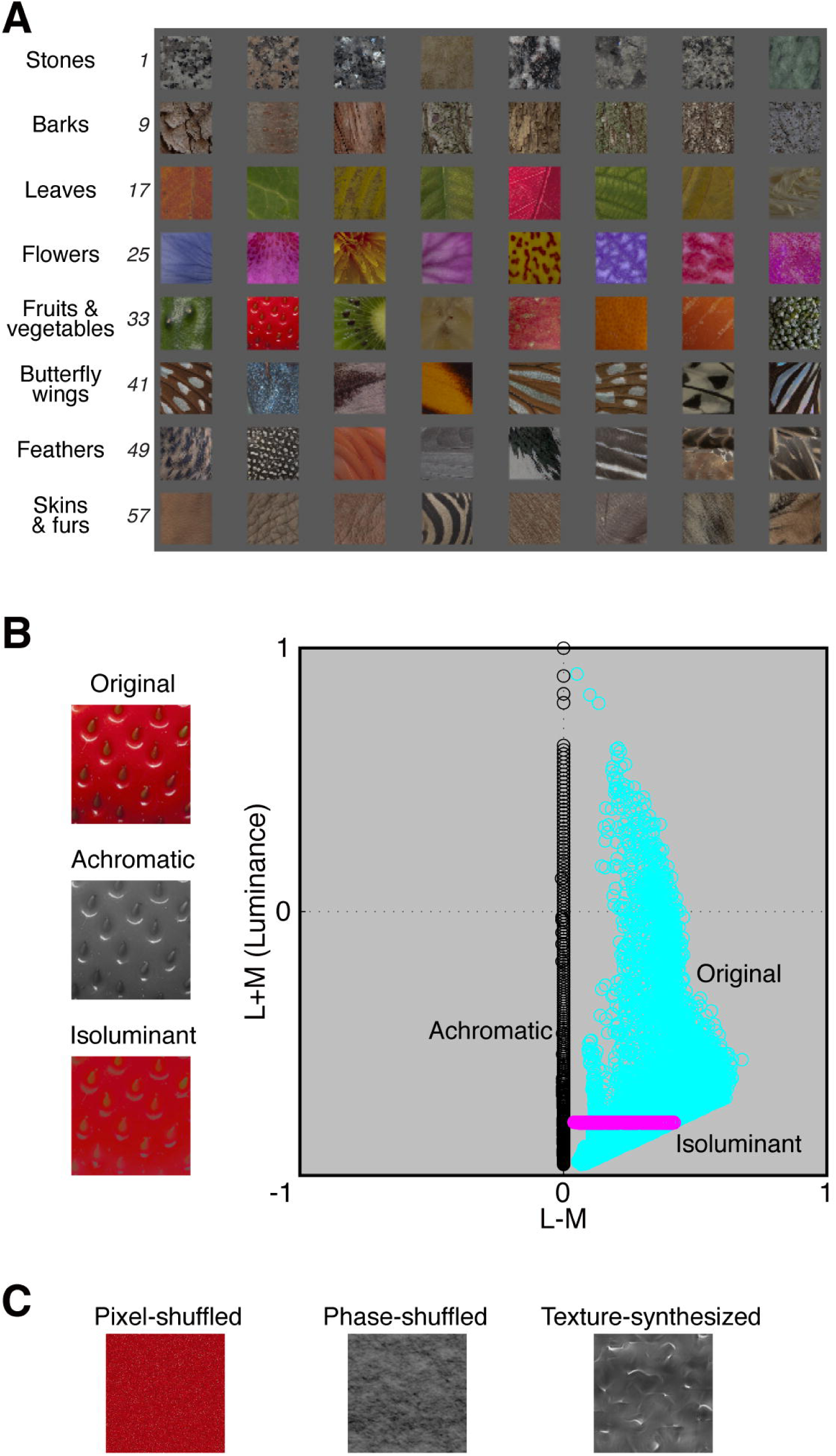
Visual stimuli. ***A***, A set of 64 surface images of natural objects (original images). We numbered stimuli consecutively from left to right and then from top to bottom. Stones (*#1–8*), tree bark (*#9–16*), leaves (*#17–24*), flowers (*#25–32*), fruits and vegetables (*#33–40*), butterfly wings (*#41–48*), feathers (*#49–56*), and skin and fur (*#57–64*). We included a blank image (*#65*) that was the same color (gray) as the background for all the images. ***B***, An example achromatic and isoluminant image. Left: The original image (top), achromatic image (middle) and isoluminant image (bottom) of a strawberry. Right: Distributions of pixel color values in DKL color space for the original (cyan), achromatic (black) and isoluminant (magenta) images. Each pixel has three values in DKL color space: L−M, S−(L+M), and L+M. In the graph, horizontal and vertical axes represent L−M and L+M, respectively. The S−(L+M) axis was not plotted for visualization purposes. ***C***, Examples of images whose spatial structures were manipulated. Shown are a pixel-shuffled (left), phase-shuffled (center), and texture-synthesized (right) image of the strawberry.

Stimulus images were adjusted, evaluated, and manipulated in DKL color space (Derrington et al. 1984; Fig. 1B). In DKL color space, one axis represents luminance corresponding to the sum of the outputs of the L- and M-cones (L+M axis), and the other two axes represent the difference between the outputs from the L- and M-cones (L−M axis) and the difference between the outputs of the S cones and the sum of the outputs from the L- and M-cones (S−(L+M) axis). We set the maximum and minimum values in the DKL color space to +1 and −1, respectively, and used D65 (Commission internationale de l'éclairage; CIE) as the white point, which was the origin of the coordinate axes of the DKL color space. The average luminance of the stimulus images and the gray background was set to −0.8 of the L+M axis and was 12 cd/m^2^.

We quantified each image with 31 image-related measures. We obtained 12 image statistics, one from each of 12 sub-band images (12 filters: three spatial frequencies [8, 16, 32 cycles/image] × four orientations [0°, 45°, 90°, 135°]) using a wavelet transform (Heeger and Bergen 1995; Portilla and Simoncelli 2000). Images were transformed to gray scale in the DKL color space before the wavelet transform. For the output values from each filter, we calculated the sum of squares for the values and used their logarithms as a measure. We obtained another seven image statistics from pixel histograms: the standard deviation (SD), skewness, and kurtosis of luminance (L+M) values across all the pixels, and the mean and SD across all the pixels’ L−M and S−(L+M) values in the DKL color space. We did not use the mean luminance of the images, because we set it to be the same across images. By applying a two-dimensional difference of Gaussian filters, we obtained the other 12 image statistics from local color contrasts (12 filters: four center-surround color combinations [L−M, M−L, S−(L+M), (L+M)−S] × three center-filter diameters [1/16, 1/32, 1/64 of image]; the size of the surround-filters was 1.6 times the size of the center-filter). For the output values from each filter, we calculated the sum of squares of the positive values and used their logarithms as a measure. We generated five types of manipulated images. Achromatic images were generated by setting the L−M and S−(L+M) values of all the pixels to zero (Fig. 1B). Photometrically isoluminant images were generated by setting the L+M (luminance) values of all pixels equal to each other and to the background (*i.e.*, −0.8; Fig. 1B). Because of the display limitation, L−M and S−(L+M) values were adjusted to conform to the display if necessary. Pixel-shuffled images

(Fig. 1C) were generated by randomly shuffling the x-y position of each pixel. Thus, the pixel histograms of pixel-shuffled images were equivalent to those of the corresponding original images, but their spatial structures differed. We generated phase-shuffled images and texture-synthesized images from achromatic images for technical reasons. Phase-shuffled images (Fig. 1C) were generated by randomly shuffling the phase of all spatial frequencies and all orientations of achromatic images. The spatial frequency and orientation contained in phase-shuffled images were the same as those in the corresponding achromatic images. Texture-synthesized images (Fig. 1C) were generated with the method developed by Portilla and Simoncelli (2000) using the MATLAB (MathWorks, MA, USA) toolbox obtained from their web site (http://www.cns.nyu.edu/~lcv/texture/). With this method, we obtained images that retained the V1-like filter outputs as well as the auto- and cross-correlation values of the filter outputs from the achromatic images, but had different inhomogeneously structured patterns (Portilla and Simoncelli 2000).

We conducted three types of recording sessions. In the first type, we presented the 64 original images to the IT (Monkeys A, B, and C) and to V4 and V1 (Monkeys A and C). In the second type, we presented original images, achromatic images, isoluminant images, and pixel-shuffled images to the IT (Monkeys A, B, and C) and to V4 and V1 (Monkeys A and C). In the third type, we presented achromatic images, phase-shuffled images, and texture-synthesized images to the IT and V1 (Monkeys D and F) and to V4 (Monkeys E and F).

We presented each visual stimulus for 0.2 s monocularly against a homogeneous gray background to the eye contralateral to the recording hemisphere and presented the same homogeneous gray during the 0.2-s intervals between stimulus presentations. The other eye was closed. For IT neurons, we presented stimuli at the fovea because the receptive fields (RFs) of IT neurons include the fovea and respond well to the stimuli located there (Gross et al. 1972). For V4 and V1 neurons, we presented stimuli at the center of the RF that was determined by audio-monitoring multiunit-activity while stimulating with hand-held circular disks, bars, or grating patterns. We presented images on a liquid crystal display (CG275W, Eizo, Ishikawa, Japan), which was regularly calibrated with the internal calibrator and checked with a spectrometer (Minolta CS-1000, Tokyo, Japan). The luminance values of the white and black areas were 125 cd/m^2^ and 1.3 cd/m^2^, respectively. We recorded stimulus onset and offset using a photodiode attached to the monitor. We repeated 25 or 30 blocks during each recording session with the stimuli order of each block pseudo-randomized such that each stimulus was presented once.

### Data analysis

We analyzed the responses of all the recorded and isolated neurons. We computed the magnitude of a visually evoked response to a given image stimulus as a change in firing rate by subtracting the spontaneous firing rate during the 0.1-s period immediately preceding the stimulus from the raw firing rate during the 0.2-s stimulus presentation period. We shifted the start of the window for the 0.2-s stimulus presentation period to 80 ms after stimulus onset for IT neurons to compensate for response latency. We evaluated the responsiveness of each neuron by comparing the firing rates during the stimulus presentation period across stimuli (*P* < 0.01, Kruskal–Wallis test). We determined the statistical significance of a response by comparing the firing rates during visual stimulation with the firing rates during the 0.1-s period immediately preceding stimulus (*P* < 0.01, Wilcoxon signed-rank test).

We evaluated the stimulus selectivity of neurons with a sparseness index (Rolls and Tovee 1995; Vinje and Gallant 2000; Tamura et al. 2014). We calculated the index with the following formula:

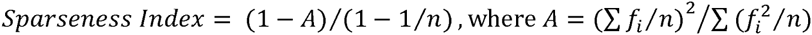

and is the activity fraction, *f*_*i*_ is the change in firing rate from the spontaneous firing rate during the stimulus presentation period of the i-th stimulus. *f*_*i*_ is the average across trials. Negative *f*_*i*_ was set to zero. *n* is the number of stimuli. The sparseness index has a range from zero to one. If a neuron responded to a smaller number of stimuli with higher firing rates, its sparseness index was closer to 1. If it responded to almost all stimuli with similar firing rates, its sparseness index was closer to 0.

We assessed a neuron’s similarity in stimulus preference between two sets of responses using Pearson’s correlation coefficient. When performing the statistical test for correlation coefficients, the coefficients were Fisher transformed.

## Results

### Responses of IT, V4, and V1 neurons to surface images of natural objects

We recorded the responses of 610 neurons from IT cortex (168 from Monkey A, 145 from Monkey B, and 297 from Monkey C) to the 64 images of natural object surfaces. We evaluated the responsiveness of each neuron by comparing the firing rates during the stimulus-presentation period across stimuli (*P* < 0.01, Kruskal–Wallis test). Forty-three percent of these neurons (265/610) were visually responsive (Fig.2). We found responsive neurons in all electrode penetrations, meaning that they were distributed throughout the lateral gyrus of IT cortex.

**Figure 2.**
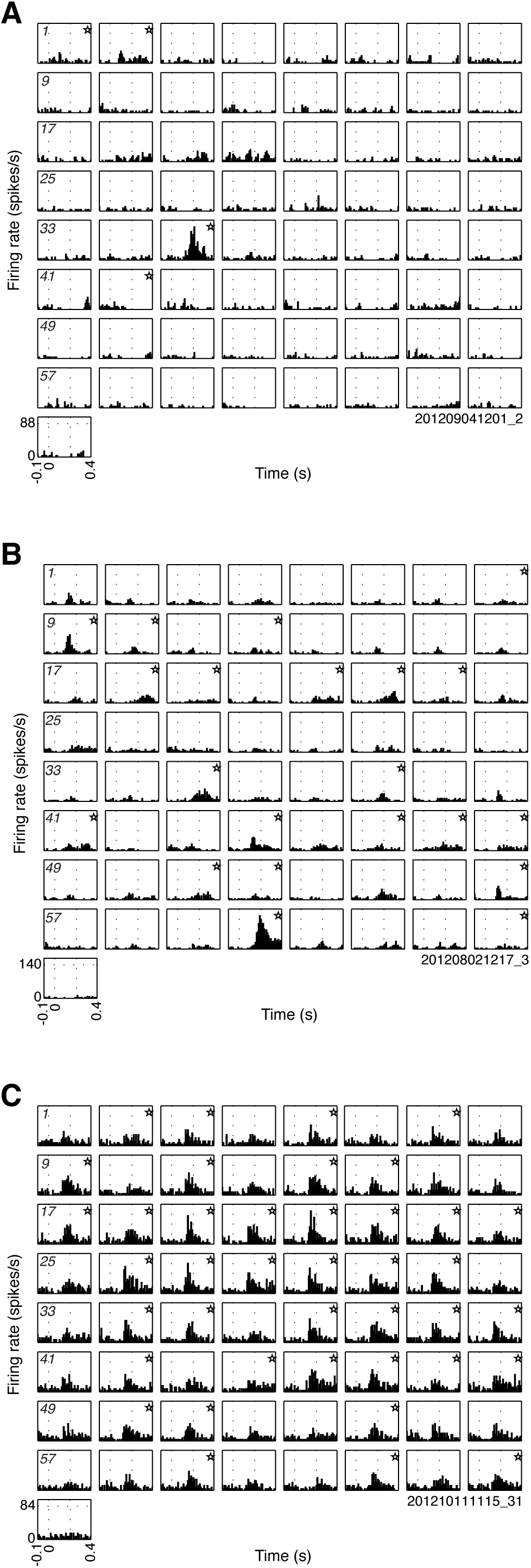
Responses of IT neurons to 64 surface images of natural objects. ***A–C***, Peristimulus time histograms (PSTHs) showing the responses of three neurons. PSTHs are arranged in the same order as the stimulus images in Figure 1A and the last PSTH (the bottom left) is for the blank image. Numbers in italic in eight leftmost PSTHs correspond to the stimulus numbers in Figure 1A. We plotted firing rates (spikes/s) for 0.5-s periods with a 10-ms bin-width. The vertical dotted lines within each PSTH indicate stimulus onset (0 s) and offset (0.2 s). Stars in the PSTHs indicate significant differences between stimulation-induced firing rates and spontaneous firing rates (*P* < 0.01, Wilcoxon signed-rank test).

Many single IT neurons responded sparsely to the set of 64 surface images. The neuron in Figure 2A was visually responsive (*P* < 0.01, Kruskal–Wallis test). It responded to four stimuli (*P* < 0.01, Wilcoxon signed-rank test) and only one stimulus (*#35*) evoked strong responses. We quantified stimulus selectivity using a sparseness index that ranges from zero to one. Neurons that respond with high firing rates to a small number of stimuli have higher sparseness indices (i.e., very selective), whereas those that respond with similar firing rates to almost all stimuli have very low indices (i.e., non-selective). The sparseness index of the neuron in Figure 2A was 0.91. The stimulus selectivity of the neuron in Figure 2B was also sharp, but broader than that of the neuron in Figure 2A (sparseness index, 0.67). For comparison, the neuron in Figure 2C had much broader stimulus selectivity (sparseness index, 0.20).

Neurons in V4 and V1 also responded to the set of 64 surface images. We recorded 578 neurons from V4 (222 from Monkey A and 356 from Monkey C) and 890 neurons from V1 (543 from Monkey A and 347 from Monkey C) for this analysis. Among the recorded neurons, 67% of V4 neurons (387/578) and 53% of V1 neurons (474/890) were visually responsive to the images (*P* < 0.01, Kruskal–Wallis test; Fig. 3A). The percent of visually responsive neurons differed among the three cortical areas (*P* < 0.001, *χ*^2^-test), with IT cortex having the lowest incidence of responsive and V4 having the highest. Sparseness indices also differed among the three cortical areas (median sparseness index: IT, 0.74; V4, 0.75; V1, 0.66; *P* < 0.001, Kruskal–Wallis test; Fig. 3B). While the sparseness index of IT neurons was similar to that of V4 neurons, it was larger than that of V1 neurons. This means that stimulus selectivity was much sharper in IT and V4 neurons than in V1 neurons.

**Figure 3.**
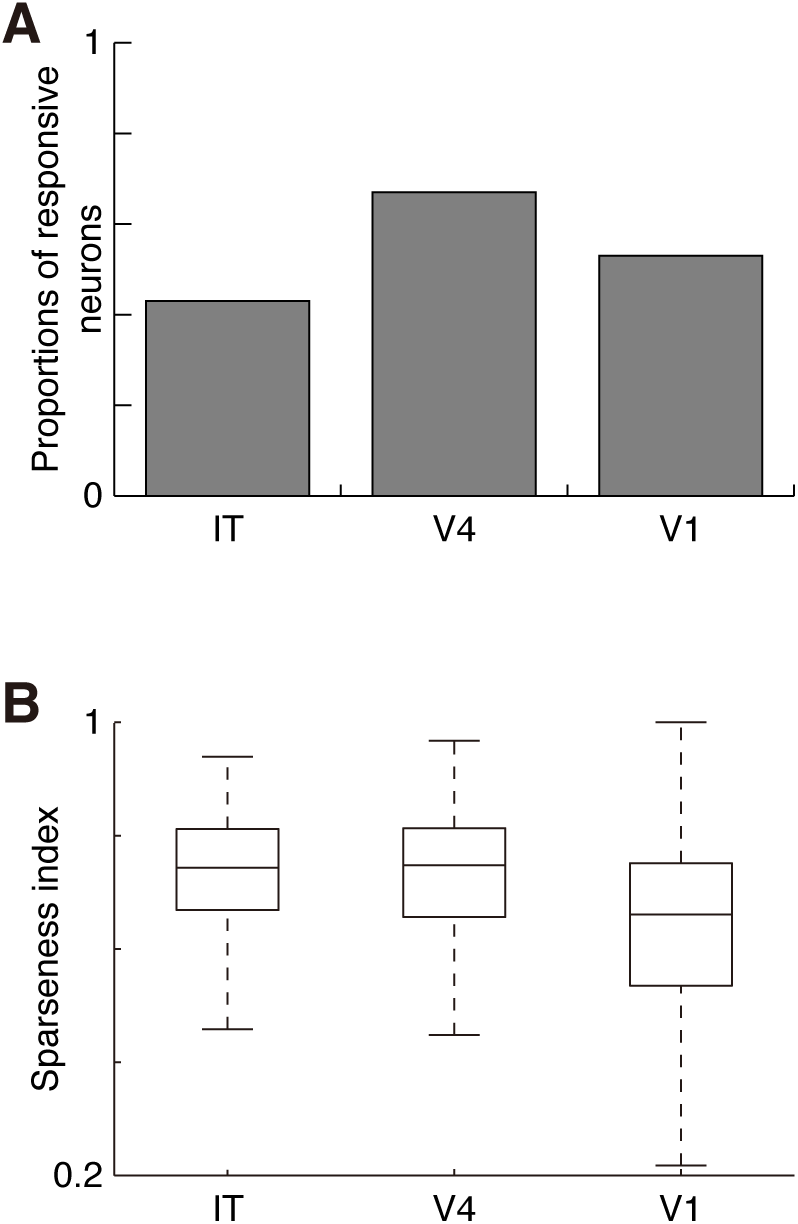
Comparisons of response properties among IT, V4, and V1 neurons. ***A***, Proportions of visually responsive neurons to the surface images of natural objects (IT, n = 610; V4, n = 578; V1, n = 890). We evaluated the responsiveness of each neuron by comparing the firing rates during the stimulus presentation period across stimuli (*P* < 0.01, Kruskal–Wallis test). ***B***, Box-plot comparison of the sparseness index. Analysis was conducted with visually responsive neurons (IT, n = 265; V4, n = 387; V1, n = 474). The center of each box is the median, and the top and bottom of the box are the upper and lower quartiles, respectively. The attached whiskers connect the most extreme values within 150% of the interquartile range from the end of each box.

Responses of IT neurons to the surface images of natural objects were independent from low-level image properties, such as distributions of pixel values, orientation, spatial frequency, or color contrasts. We quantified each stimulus image with 31 image statistics (seven statistics from pixel histograms, 12 from sub-band images, and 12 from local color contrasts; see Materials and Methods for details). We calculated *r*^2^-values between a neuron’s 64 responses (a vector with 64 elements) and one of the statistics (another vector with 64 elements). By repeating this process for each of the 31 statistics, we obtained 31 *r*^2^-values for each neuron, and designated a neuron’s largest *r*^2^-value as its representative of *r*^2^-value. We found that only a small fraction of IT neurons (12.8%, 34/265) showed significant correlation between the image statistics and responses. The median *r*^2^-value for IT neurons was 0.09 (*n* = 265; Fig. 4A and B), meaning that 9% of responses could be explained with the image statistics. Responses of most V4 neurons were also nearly independent from the image statistics (median of *r*^2^-value = 0.08; *n* = 387; Fig. 4B). In contrast, responses of V1 neurons were weakly related to the image statistics (median = 0.15; *n* = 474; Fig. 4B). The *r*^2^-values differed among the three cortical areas (*P* < 0.001, Kruskal–Wallis test). Thus the simple image statistics explained at least a part of V1 neuronal responses to the surface images of natural objects, whereas they failed to explain the responses of most IT and V4 neurons.

**Figure 4.**
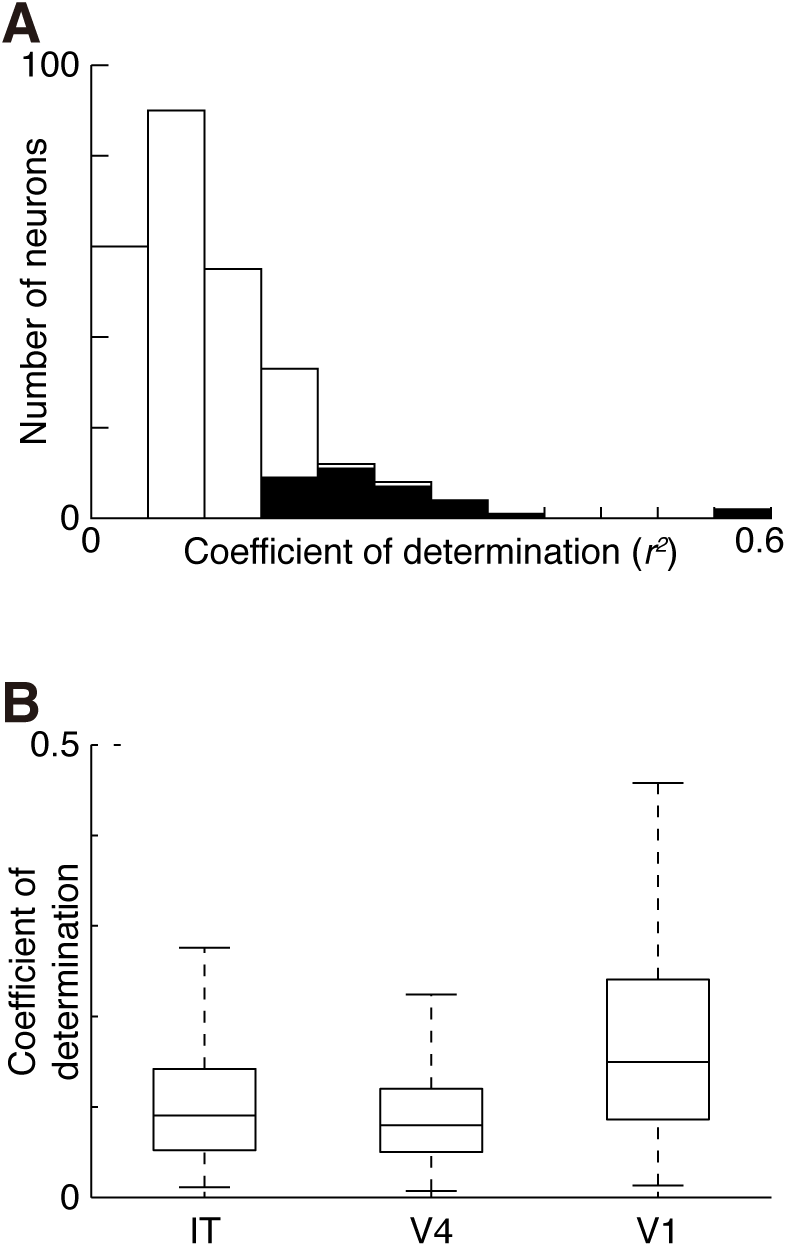
Relationship between responses and image statistics in IT, V4, and V1 neurons. ***A***, Frequency distributions of the coefficients of determination (*r*^2^) for the IT neurons. We calculated *r*^2^-values between responses and each of the 31 image statistics and plotted the largest among the 31 *r*^2^-values for each neuron. Filled columns, *r*^2^-values significantly larger than those calculated with shuffled responses (*P* < 0.01, permutation test, 1,000 times of randomization). Open columns, non-significant *r*^2^-values. ***B***, Comparison of *r*^2^-values among IT (n = 265), V4 (n = 387), and V1 (n = 474) neurons.

### Responses of IT, V4, and V1 neurons to manipulated surface images of natural objects

Surface images of natural objects contain a variety of visual features, such as color, luminance-contrast, and spatial structures. To gauge the contribution that each of these features had on the responsiveness and stimulus preference of neurons exposed to natural surface images, we recorded responses of single neurons to the original images, achromatic images (Fig. 1B), isoluminant images (Fig. 1B) and pixel-shuffled images (Fig. 1C, left), respectively. We examined responses to these manipulated images with another set of 107 neurons from IT (32 from Monkey A, 25 from Monkey B, and 50 from Monkey C), 127 neurons from V4 (84 from Monkey A and 43 from Monkey C), and 50 neurons from V1 (16 from Monkey A and 34 from Monkey C). To examine how the stimulus manipulations affect responses, we selected those (35 in IT, 79 in V4 and 34 in V1) that were responsive to the original images (*P* < 0.01, Kruskal–Wallis test) and showed significant response to the best original image (the one that induced the largest response, *P* < 0.01, Wilcoxon signed-rank test).

#### Responses to achromatic images

A small but significant fraction of IT neurons (34%, 12/35) maintained visual responsiveness to the achromatic version of the best original images (Fig. 5A–C, black lines and circles; Fig. 6A). The response magnitudes of IT neurons to the achromatic version were smaller than those to the best original images (*P* < 0.001, Wilcoxon signed-rank test; Fig. 5A and B), and the mean response magnitude was 0.37 ± 0.40 (n = 35; Fig. 6A, middle) when normalized to the responses to the best original images. Thus, colors augmented, or were crucial, for responses of IT neurons to surface images of natural objects.

**Figure 5.**
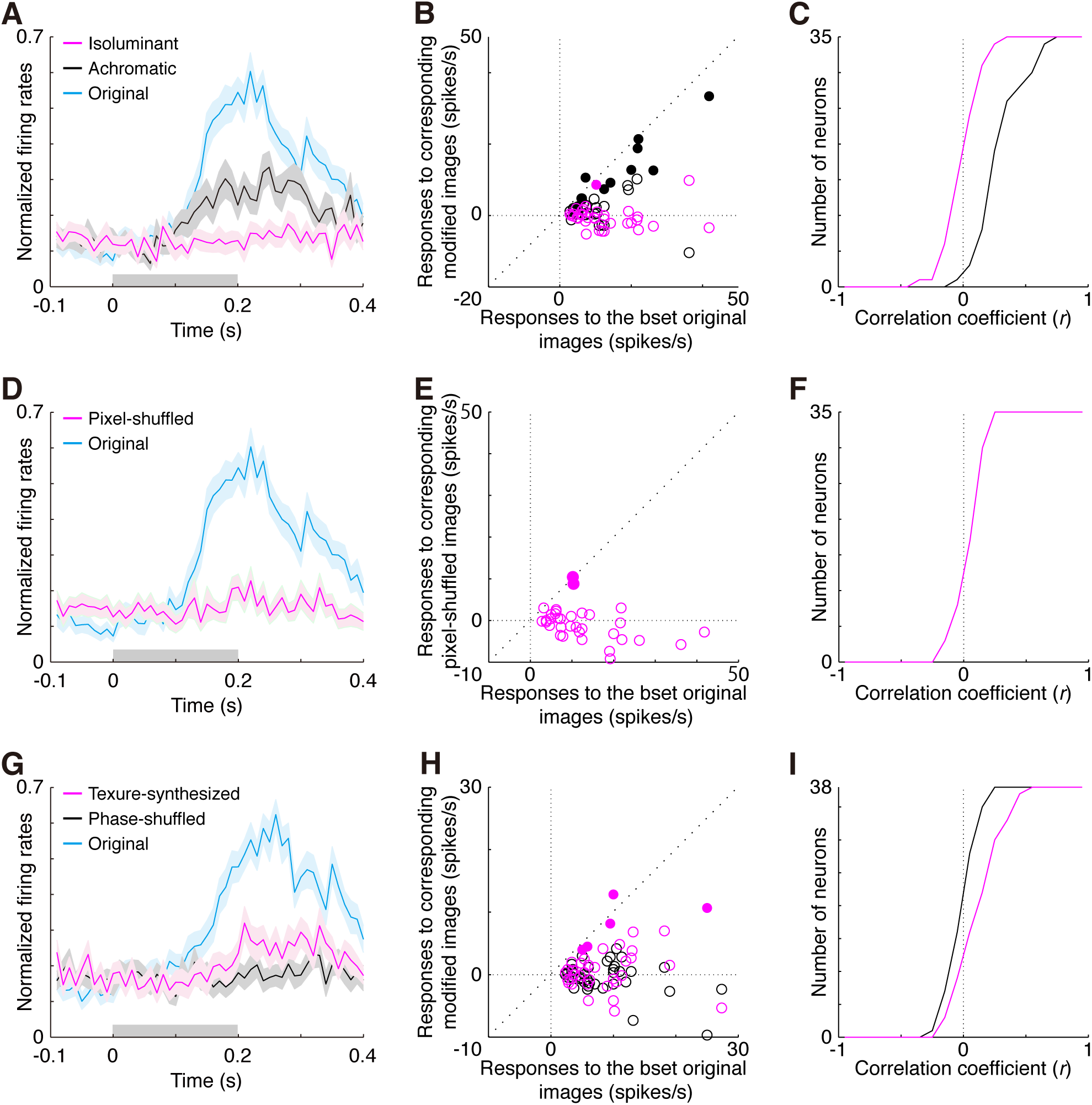
Responses of IT neurons to manipulated surface images of natural objects. ***A***, Population PSTHs for original images (cyan), achromatic images (black), and isoluminant images (magenta). We constructed population PSTHs in the following way. For each neuron, we determined the best original image (the one that induced the largest response), and normalized the PSTHs with the peak-firing rates. We obtained population PSTHs by averaging the normalized PSTHs across neurons. We constructed population PSTHs for the achromatic or isoluminant version of the best original images in the same way but normalized by the peak-firing rates in the PSTH of the best *original* image. Shades around the line represent the standard error of the mean. The gray bar on the horizontal axis indicates the stimulus presentation period (0–0.2 s). ***B***, Scatter plot of response magnitudes for the best original images and the achromatic images (black circles) and for the best original images and the isoluminant images (magenta circles). The response magnitudes (firing rates) for the best original images are plotted on the horizontal axis, and those for the corresponding manipulated images are plotted on the vertical axis. Each circle represents a response pair from a neuron. Filled and open circles represent statistically significant and non-significant responses to the manipulated images, respectively. We determined the statistical significance of a response by comparing the firing rates during visual stimulation with the firing rates during the 0.1-s period immediately preceding stimulus presentation (*P* < 0.01, Wilcoxon signed-rank test). Note that all responses to the best original images were statistically significant. The diagonal broken line is the identity line. ***C***, Cumulative frequency distribution of the correlation coefficients between responses to the original images and those to the achromatic images (black) or the isoluminant images (magenta). ***D***, Population PSTHs for the best original images (cyan) and the corresponding pixel-shuffled images (magenta). ***E***, Comparison between response magnitudes for the best original images and for the pixel-shuffled images. ***F***, Cumulative frequency distributions of the correlation coefficients between responses to the original images and to the pixel-shuffled images. ***G***, Population PSTHs for the best achromatic images (cyan), the corresponding phase-shuffled images (black), and the corresponding texture-synthesized images (magenta). ***H***, Comparison between response magnitudes for the best achromatic images and the phase-shuffled images (black circles), or the texture-synthesized images (magenta circles). ***I***, Cumulative frequency distribution of the correlation coefficients between responses to the achromatic images and the phase-shuffled images (black), or the texture-synthesized images (magenta). Note that phase-shuffled images and texture-synthesized images were achromatic. We analyzed 35 IT neurons for A–F and another 38 IT neurons for G–I.

**Figure 6.**
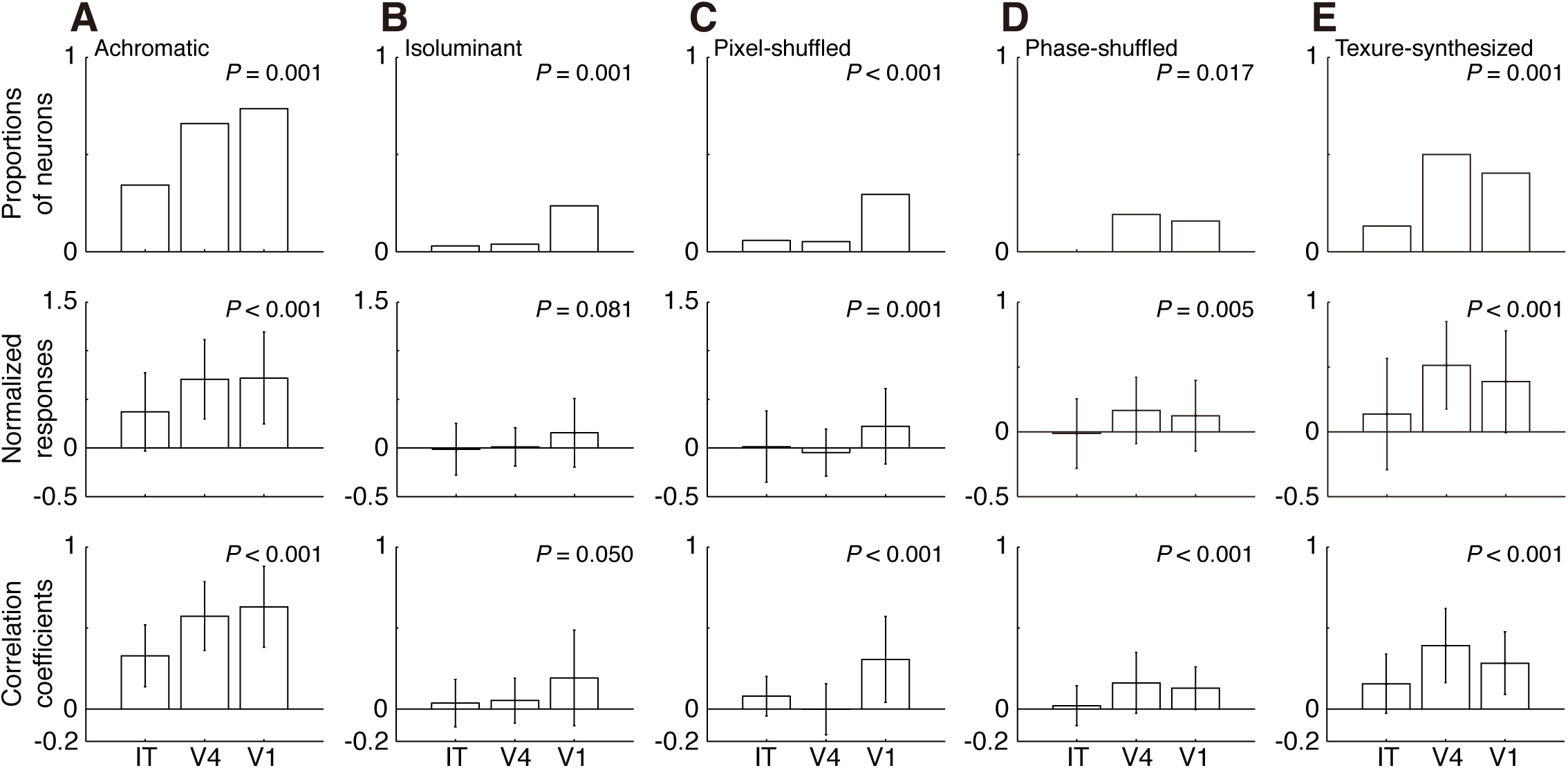
Responses to manipulated images in IT, V4, and V1 neurons. Among neurons responsive to the original images, we compared proportions of neurons responsive to manipulated images (Top), response magnitudes to manipulated images normalized with those to the best original images (middle), and correlation coefficients between responses to a set of manipulated images and those to the original images (bottom). ***A–C***, Comparisons of respective responses to achromatic, isoluminant, and pixel-shuffled images across the three cortical areas. For A, B and C, we analyzed 35 IT, 79 V4, and 34 V1 neurons. ***D–E***, Comparisons of respective responses to phase-shuffled and texture-synthesized images across the three cortical areas. For D and E, we analyzed 38 IT, 84 V4, and 57 V1 neurons. Error bars for normalized responses (middle row) and correlation coefficients (bottom row) are standard deviations. *P*-values from statistical comparisons among IT, V4, and V1 (*χ*^2^-test for proportion of visually responsive neurons, Kruskal–Wallis test for normalized response magnitudes and for correlation coefficients) are provided in each panel.

To evaluate contributions of color to a neuron’s stimulus preference, we examined the similarity in stimulus preference between the original and achromatic images by calculating correlation coefficients between the responses associated with each image type. If the two sets of responses are similar to each other (*i.e.*, a correlation coefficient close to 1), color does not contribute to the stimulus preference. Analysis showed that the mean correlation coefficient was 0.33 ± 0.19 (n = 35) and was different from zero (*P* < 0.001, Wilcoxon signed-rank test; Fig. 5C, black line; Fig. 6A, bottom), meaning that stimulus preferences of IT neurons evaluated with achromatic images were similar in some way to those evaluated with original images. The results thus suggest that some part of the stimulus preference exhibited by IT neurons to surface images of natural objects can be maintained without color information.

Most V4 and V1 neurons responded at least weakly to achromatic images, and maintained their stimulus preferences (Fig. 6A). The proportion of neurons, the normalized response magnitudes, and the similarity in stimulus preference differed among IT, V4, and V1 neurons (proportion of neurons, *P* = 0.001, *χ*^2^-test; normalized response magnitudes, *P* < 0.001, Kruskal–Wallis test; similarity in stimulus preferences, *P* < 0.001, Kruskal–Wallis test; Fig. 6A). Specifically, IT neurons showed smaller proportion of responsive neurons, smaller response magnitudes, and less correlation than V4 and V1 neurons. Thus, the color contributes to surface-image responses in IT more than in V4 or V1.

#### Responses to isoluminant images

Almost all IT neurons lost visual responsiveness to the isoluminant version of the best original images (Fig. 5A–C, magenta lines and circles; Fig. 6B), and only one neuron maintaining its visual responsiveness (1/35, 3%; Fig. 6B, top). The response magnitudes were also smaller than those for the original images (*P* < 0.001, Wilcoxon signed-rank test; Fig. 5A and B), and the mean normalized response magnitude was less than zero (−0.14 ± 0.27 Fig. 6B, middle). The mean correlation coefficient between responses to the two types of images was not different from zero (0.04 ± 0.15; *P* = 0.219, Wilcoxon signed-rank test; Fig. 5C, magenta line; Fig. 6B, bottom). This can be explained by the unresponsiveness to isoluminant images. Thus, luminance contrast, but not color contrast, was essential for IT neuronal responses to surface images of natural objects.

A similar situation was observed in V4, with only a few neurons maintaining responsiveness to isoluminant images. In contrast, a small but much larger proportion of V1 neurons did respond to isoluminant images (Fig. 6B). Comparing these results across regions showed that the proportion of visually responsive neurons to isoluminant images differed among IT, V4, and V1 (*P* = 0.001, *χ*^2^-test; Fig. 6B, top), with IT and V4 neurons showing smaller proportions than V1. The normalized response magnitude and the correlation coefficients between responses to the original images and isoluminant images did not differ among the three cortical areas (normalized response magnitude, *P* = 0.081, Kruskal–Wallis test, Fig. 6B, middle; correlation coefficients, *P* = 0.050, Kruskal–Wallis test, Fig. 6B, bottom). Overall, the removal of luminance contrast from the surface images of natural objects significantly diminished the responses of neurons in all the three cortical areas, and the effect was much stronger for IT and V4 neurons.

#### Responses to pixel-shuffled images

Only two IT neurons (6%, 2/35) maintained their responsiveness to the pixel-shuffled version of the best original images (Fig. 5D–F, magenta lines and circles; Fig. 6C). The response magnitudes of IT neurons to pixel-shuffled images were smaller than those to the best original images (*P* < 0.001, Wilcoxon signed-rank test; Fig. 5D and E), and the mean normalized response magnitude was almost zero (0.01 ± 0.37; Fig. 6C, middle). The mean correlation coefficient between responses to the original images and pixel-shuffled images was almost zero (0.08 ± 0.12; Fig. 5F; Fig. 6C, bottom), although it was statistically greater than zero (*P* = 0.001).

While most V4 neurons also lost visual responsiveness to pixel-shuffled images, a large fraction of V1 neurons responded to pixel-shuffled images and maintained stimulus preferences (Fig. 6C). The proportion of responsive neurons and the normalized response magnitude to pixel-shuffled images differed among IT, V4, and V1 (proportion of neurons, *P* < 0.001, *χ*^2^-test; response magnitudes, *P* = 0.001, Kruskal–Wallis test; Fig. 6C, top, middle). The correlation coefficient between responses to the original images and pixel-shuffled images also differed among brain regions (*P* < 0.001, Kruskal–Wallis test; Fig. 6C, bottom). Pixel-wise spatial structures of the surface images of natural objects were essential for responses of most IT and V4 neurons. In contrast, the shape of pixel histograms, which was maintained in pixel-shuffled images, could explain a part of V1 neuronal responses.

#### Responses to phase-shuffled images

To further evaluate contributions of spatial structures, we examined responses of neurons to phase-shuffled and texture-synthesized images. Because phase-shuffled and texture-synthesized images were generated from achromatic images (for technical reasons), we compared responses between the best achromatic images and their corresponding phase-shuffled and texture-synthesized images. Phase-shuffled images retained spatial frequency and orientation components of achromatic images, but had different spatial phases (Fig. 1C, center). Texture-synthesized images retained V1-like filter outputs and local second-order image statistics, but had different inhomogeneously structured patterns (Fig. 1C, right).

We recorded responses from 259 IT neurons (65 from Monkey D and 194 from Monkey F), 165 V4 neurons (38 from Monkey E and 127 from Monkey F) and 91 V1 neurons (22 from Monkey D and 69 from Monkey F). We restricted analyses to 38 neurons in IT, 84 neurons in V4, and 57 neurons in V1 that were responsive to achromatic images (*P* < 0.01, Kruskal–Wallis test) and showed significant response to the best achromatic image (the one that induced the largest response, *P* < 0.01, Wilcoxon signed-rank test).

All IT neurons lost responsiveness to phase-shuffled version of the best achromatic images (Fig. 5G–I, black lines and circles; Fig. 6D), and the mean normalized response magnitude was almost zero (−0.01 ± 0.27; Fig. 6D, middle). The mean correlation coefficient between responses to achromatic images and phase-shuffled images was not different from zero (0.02 ± 0.12; *P* = 0.357, Wilcoxon signed-rank test; Fig. 5I, black line; Fig. 6D, bottom), as expected from the unresponsiveness.

A small number of V4 and V1 neurons maintained visual responsiveness and preference to phase-shuffled images (Fig. 6D), and their proportions did not differ among IT, V4, and V1 (*P* = 0.017, *χ*^2^-test; Fig. 6D, top). However, the mean normalized response magnitude to phase-shuffled images (*P* = 0.005, Kruskal–Wallis test; Fig. 6D, middle) and the correlation coefficient between responses to achromatic images and phase-shuffled images (*P* < 0.001, Kruskal–Wallis test; Fig. 6D, bottom) did differ across regions. Thus, the importance of spatial phase was larger in IT than in V4 and V1.

#### Responses to texture-synthesized images

A small fraction (13%, 5/38) of IT neurons maintained responsiveness to the texture-synthesized version of the best achromatic images whose inhomogeneously structured patterns were different from the original images (Fig. 5G–I, magenta lines and circles; Fig. 6E). The response magnitudes were smaller than those to achromatic images (*P* < 0.001, Wilcoxon signed-rank test; Fig. 5G and H), and the mean normalized response magnitude was 0.14 ± 0.43 (Fig. 6E, middle). The correlation coefficient between responses to achromatic images and texture-synthesized images was 0.16 ± 0.18, which was different from zero (*P* < 0.001; Fig. 5I, magenta line; Fig. 6E, bottom). Presence of neurons responsive to texture-synthesized images suggests the importance of spatial structures captured in the V1-like filter outputs and local second-order image statistics for the responses of IT neurons. At the same time, since many neurons decreased responsiveness, inhomogeneously structured patterns, which were modified in the texture-synthesized images from the original, were important for the responses of IT neurons to surface images of natural objects.

Much larger fractions of V4 and V1 neurons maintained visual responsiveness and stimulus preference to texture-synthesized images. The proportions of responsive neurons and the normalized response magnitude to texture-synthesized images differed among IT, V4, and V1 (proportion of neurons, *P* = 0.001, *χ*^2^-test; Fig. 6E, top; response magnitudes, *P* < 0.001, Kruskal–Wallis test; Fig. 6E, middle). The correlation coefficients between responses to achromatic images and texture-synthesized images also differed across regions (*P* < 0.001, Kruskal–Wallis test; Fig. 6E, bottom). Thus, contributions of spatial structures that can be represented with local second-order image statistics were larger for V4 and V1 neuronal responses to surface images of natural objects than they were for IT neurons.

### Relationships between responses to different types of manipulated surface images

Finally, we examined how the effect of one type of image manipulation related to that of another type of manipulation. If visual features themselves or their neural representations are related to each other, the effects of different image manipulations should also be related. We evaluated these relationships by correlating the responses of neurons to manipulated images of different types. We presented three types of manipulated images to one set of neurons, resulting in three pairs for comparison, and two types of manipulated images to another set of neurons, resulting in a fourth pair for analysis.

In IT, responses to isoluminant images correlated positively with those to pixel-shuffled images (*r* = 0.47, *P* = 0.005; Fig. 7A, top-right). These results indicated that neurons that received luminance contrast-based information also received pixel-wise spatial information, and vice versa. There were no significant correlations in any of the other response pairs in IT neurons (*P ≥* 0.01). In V4, responses to phase-shuffled images positively correlated with those to texture-synthesized images (*r* = 0.31, *P* = 0.004; Fig. 7B, middle), meaning that V4 neurons that responded to phase-shuffled images also responded to texture-synthesized images. There were no significant correlations in any of the other response pairs in V4 neurons (*P ≥* 0.01). Responses of V1 neurons to achromatic images were negatively correlated with those to isoluminant images (*r* = −0.57, *P* < 0.001; Fig. 7A, bottom-left), meaning that V1 neurons that were tolerant to removal of color were intolerant to removal of luminance-contrast, and vice versa. V1 responses to isoluminant images were positively correlated with those to pixel-shuffled images (*r* = 0.78, *P* < 0.001; Fig. 7A, bottom-right), meaning that V1 neurons that were tolerant to removal of luminance-contrast were also tolerant to pixel shuffling. There were no significant correlations in any of the other response pairs in V1 (*P ≥* 0.01). These data show that responses to different types of manipulated surface images did not share common relationships in the three cortical areas, thus indicating that each cortical area integrates these features in different ways.

**Figure 7.**
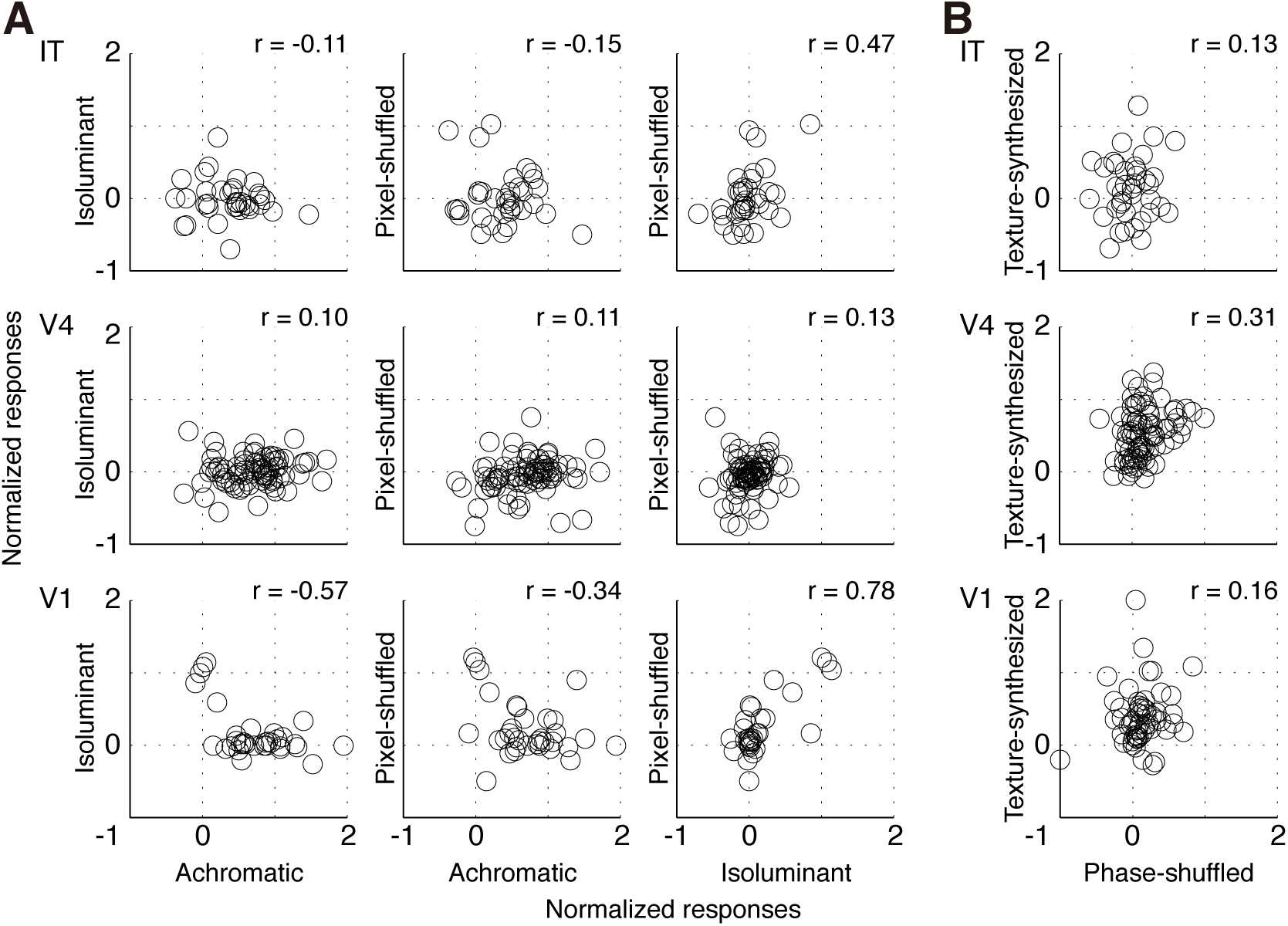
Relationships between the responses to different pairs of manipulation type for IT, V4, and V1 neurons. ***A***, Response relationship for achromatic vs. isoluminant images (left column), achromatic vs. pixel-shuffled images (middle column), and isoluminant vs. pixel-shuffled images (right column) for IT (top row, n = 35), V4 (middle row, n = 79), and V1 (bottom row, n = 34) neurons. For each neuron, we determined the most effective original image. Response magnitudes for the manipulated images were normalized to the corresponding best images. Normalized responses to two types of manipulated image are plotted against each other. Each dot represents data from one neuron. The correlation coefficient is provided in each panel. ***B***, Relationships between responses to phase-shuffled and texture-synthesized images for IT (top row, n = 38), V4 (middle row, n = 84), and V1 (bottom row, n = 57) neurons. Here, we normalized response magnitudes for manipulated images to those for the best corresponding achromatic images.

## Discussion

In the present study, we examined responses of IT neurons to images of natural object surfaces and compared them with those of V4 and V1 neurons. We found that about half of IT neurons responded to the surface images with sharp stimulus selectivity, and that their responses were independent from simple image statistics. Removal of color, removal of luminance contrasts, and degradation of spatial structures significantly decreased responses in these IT neurons. Comparing responses among IT, V4, and V1 revealed that IT and V4 neurons exhibited a similar degree of stimulus selectivity, which was sharper than that of V1 neurons. V1 neuronal responses were weakly but significantly correlated to image statistics, whereas response in IT and V4 were not. In all the three cortical areas, neural responses to modified images were significantly less than those to the original images, but the degree of reduction differed across regions. Some types of image modification reduced responses in IT more than in V4 and V1, whereas others reduced IT and V4 responses more than V1 responses.

### Responses of IT neurons to surface images of natural objects

We found that about half of IT neurons responded selectively to surface images of natural objects. Neurons in IT cortex have repeatedly been shown to respond selectively to shapes (Gross et al. 1972; Desimone et al. 1984; Tanaka et al. 1991), with some even being insensitive to surface features that define the contour (Sáry et al. 1993). Based on these studies, we expected that only a small fraction of IT neurons would respond to surface images of natural objects. However, we found that about half of all the recorded neurons in IT cortex responded to the images. We recorded responsive neurons in all electrode penetrations made in the lateral gyrus part of IT cortex. If we had recorded neurons around the superior temporal sulcus or the anterior middle temporal sulcus, where neurons responsive to surface-related features have been reported (Komatsu et al. 1992; Köteles et al. 2008), responsiveness might have been even more prevalent than we found here.

In the present study, we found that IT neurons had sharp stimulus selectivity and that their responses to surface images of natural objects were independent from simple image statistics. These characteristics can be explained through integration of visual features by single IT neurons. Indeed, consistent with this, removal or degradation of any visual features reduced or eliminated the responses of most IT neurons. Integration of simple patterns and colors by single IT neurons has been reported (Desimone et al. 1984; Tanaka et al. 1991; Komatsu and Ideura 1993). Here, we extend these findings by showing that most IT neurons that respond to surface images of natural objects require multiple surface features for their activation.

Each visual feature contributed to IT neuronal responses in a different manner. We found that removal of color reduced response magnitude, but selectivity of some IT neurons was maintained. In contrast, removing luminance contrast and degrading spatial structures eliminated responsiveness of most IT neurons. These results suggest that luminance contrast and spatial structure are essential for IT responses to surface images of natural objects, whereas color serves to augment responses. Because removing color did not alter stimulus preference, color does not seem to contribute a great deal to stimulus preference in IT neurons responsive to surface images of natural objects. Rather, stimulus preference of IT neurons to surface images of natural objects was largely determined by luminance contrast-based spatial structures.

### Comparisons of responses to surface images of natural objects among IT, V4, and V1

Response properties of IT neurons to surface images of natural objects are likely acquired via gradual and hierarchical processing in the ventral visual cortical pathway. Comparisons across regions can reveal the level at which these response properties emerge. For example, the degree of stimulus selectivity—as quantified by the sparseness index—was lower in V1 neurons than in V4 or IT neurons. Additionally, the relationship between responsiveness and image statistics was higher in V1 neurons than in V4 or IT neurons. These properties are thus modified at the level between V1 and V4. Similarly, responses to isoluminant images and pixel-shuffled images exhibited less reduction in V1 neurons than in V4 or IT neurons, indicating that the importance of luminance contrast and pixel-wise spatial structure increases between V1 and V4. Other response properties appear to be modified between V4 and IT. For example, responses to achromatic images, phase-shuffled images, and texture-synthesized images exhibited greater reduction in IT neurons than in V4 or V1 neurons, indicating that the importance of color, spatial phase, and higher order spatial structures increases between V4 and IT. Thus, selective responses of IT neurons to surface images of natural objects were constructed not simply by inheriting the response properties from the preceding areas but by gradual and hierarchical processing in the ventral pathway. Some changes in response properties along the ventral pathway have been reported (Tanaka et al. 1991; Kobatake and Tanaka 1994; Denys et al. 2004; Rust and DiCarlo 2012; Goda et al. 2014; Namima et al. 2014), and the present results revealed that information derived from the surfaces of natural objects is also processed in a hierarchical manner.

One might argue that the differences in response properties among the three cortical areas arise from differences in RF size in relation to stimulus size. Because the RFs of V1 neurons are significantly smaller than stimulus images, image statistics calculated across whole images may not accurately represent the image properties within the RF of a V1 neuron. This discrepancy might lead to underestimation of the correlation between V1 neuronal responses and the image statistics. It may also result in large reductions in responses to modified images, which are constructed by referencing the image statistics. However, we found that the correlation coefficients between V1 responses and the image statistics were not smaller than those for IT or V4 neurons (see Fig. 4B). The degree to which V1 neurons responded less to modified images was not larger than that of IT or V4 neurons. These results suggest that the discrepancy between image statistics and image properties within the small RFs of V1 neurons was not the cause of the differences in response properties among the three cortical areas, although we cannot rule out the possibility that the correlations and responses to modified images were underestimated in V1 neurons, leading us to underestimate the real differences.

### Further processing of surface visual features in IT cortex

At the end of the hierarchical processing, visual information about object surfaces may be integrated with visual information about objects contour, creating representations of objects that is invariant and robust to modification and degradation of visual inputs. In past studies, one of the authors reported that almost all IT neurons (79%–96%) were visually responsive to object images that retain both natural outer contours and surface visual features (Tamura and Tanaka 2001; Tamura et al. 2005; Tamura et al. 2014). In the present study, we tested visual responsiveness with images that lacked contours but retained surface visual features. Analysis showed a much lower incidence of responsive neurons (43%). This result suggests that a population of IT neurons require contour information for their activation, and that they may integrate contour information with surface-related information. Indeed, some IT neurons have been shown to be sensitive to both shape and color or textures (Desimone et al. 1984; Tanaka et al. 1991; Komatsu and Ideura 1993; Edwards et al. 2003; Conway et al. 2007; Köteles et al. 2008), and a human fMRI study has also indicated the possibility that shape is integrated with surface features in the fusiform gyrus (Cavina-Pratesi et al. 2010).

Alternatively, the majority of IT neurons sensitive to contour may not be sensitive to other features, and contour information might be segregated from surface-related information within IT. Some studies have reported IT neurons sensitive to shape but insensitive to surface-related properties (Komatsu and Ideura 1993; Sáry et al. 1993; Köteles et al. 2008). Human fMRI studies and neuropsychological studies have shown that the lateral occipital cortex is involved in shape processing, whereas more medial parts are involved in processing surface-related properties (see Eagleman and Goodale 2009 for a review). We should conduct more systematic analysis at the single-neuron level to clarify how contour information and surface-related information are represented in IT cortex.

## Note

Thanks to H Takada, Y Inoue, S Ishida, K Kondo, and H Kaneko for technical assistances. Four monkeys were provided by NBRP “Japanese Monkeys” through the National BioResource Project of MEXT, Japan.

